# Potassium-selective channelrhodopsins can exert hyper- or depolarizing effects in excitable cells of *Caenorhabditis elegans*, depending on experimental condition

**DOI:** 10.1101/2024.06.14.599090

**Authors:** Christiane Ruse, Marius Seidenthal, Linda Tillert, Johannes Vierock, Alexander Gottschalk

## Abstract

One of the most frequent applications of optogenetic tools is for depolarization and stimulation of excitable cells such as neurons and muscles. Equally important, but less frequently used, are inhibitory tools that suppress activity through cellular hyperpolarization. These tools often rely on chloride conductance. Yet, *in vivo*, re- and hyperpolarization is typically mediated by potassium. In recent years, light-gated ion channels with a high preference for potassium were identified (Kalium channelrhodopsins, KCRs), and their inhibitory potential described in different organisms. Here, we characterized *Hc*KCR1 and WiChR, in cholinergic neurons and muscles of *Caenorhabditis elegans*. Hyperpolarization of these cell types both induces muscle relaxation and, consequently, an elongation of the animals. Thus, we analyzed body length before, during, and after illumination, to assess KCR effectiveness, and to benchmark stimulation parameters like light intensity and duration. For *Hc*KCR1 in cholinergic neurons, continuous illumination at high light intensities (1-4.5 mW/mm^2^) evoked only a transient elongation, while stimulation at 0.1 mW/mm^2^ could maintain inhibition for the duration of the stimulus in some transgenic strains. For animals expressing WiChR in body wall muscle cells or cholinergic neurons, we again observed brief hyperpolarization during continuous illumination, however, still during the stimulus, this changed to body contraction, corresponding to depolarization. This effect was long lasting, and required dozens of seconds for reversion, but could be reduced by pulsed illumination and fully avoided by less efficient channel activation using green or orange light. Hence, KCRs can be applied to hyperpolarize *C. elegans* cells, but require optimized illumination parameters.

**Article summary:** To inhibit excitable cells, light-gated, potassium-selective channels (KCRs) can be used. This study explores whether stimulation of KCRs *Hc*KCR1 and WiChR in cholinergic neurons and muscle cells of *Caenorhabditis elegans* can induce inhibition during illumination. While inhibition could be achieved, depending on light conditions, the authors unexpectedly also observed excitation. These effects may occur due to a combination of high conductivity of KCRs, and partial conductance of other cations. These findings highlight the need for specific experimental conditions in future studies utilizing these tools. The authors also present conditions that can partially or fully avoid the unwanted depolarizing effects.

## Introduction

Neuronal excitation or inhibition mediated through light-activated ion channels is one of the most common applications of optogenetics. Channelrhodopsin-2 (ChR2) from the green alga *C. reinhardtii*, is likely the most widely known used tool. ChR2 is a largely unselective cation channel that is activated by blue light and has been applied in many experimental systems including *C. elegans*, to stimulate excitable cells by inducing depolarization (Nagel et al. 2003; Nagel et al. 2005). Regarding inhibition, anion-conducting ChRs (ACRs) with high chloride conductance have been described and used as hyperpolarizing tools (Govorunova et al. 2015; Bergs et al. 2018). However, while chloride ions are important for setting the resting membrane potential, it is largely the role of potassium (K^+^) to mediate re- and hyperpolarization in neurons and muscles. Thus, light-regulated K^+^-channels have been a target of research for a comparably long time. For example, existing K^+^-channels were made light-responsive by chemical modification with photoswitchable agents or by genetic fusion of light-reactive domains (Banghart et al. 2004; Cosentino et al. 2015). In 2022, natural K^+^-selective ChRs (KCRs) from *Hyphochytrium catenoides*, named *Hc*KCRs, were described (Govorunova et al. 2022a) with a novel type of K^+^ selectivity filter (Govorunova et al. 2022b; Vierock et al. 2022). The selectivity filter of these channels was found to be mainly localized in the extracellular part, resulting in a higher selectivity for ions that are moving into the cells than those moving out. Consequently, these channels have displayed a lowered preference for Na^+^ over K^+^ when inverted ion gradients were applied *in vitro*, meaning high intracellular Na^+^ and high extracellular K^+^ concentrations (Tajima et al. 2023). Of the identified channels, the green light-sensitive *Hc*KCR1, appeared to have a higher specificity towards K^+^ compared to the blue light-responsive *Hc*KCR2 (Govorunova et al. 2022a). Electrophysiological measurements performed in a later study (Vierock et al. 2022) have confirmed the high K^+^-conductance but have noted a residual Na^+^-conductivity, and a transition of K^+^-efflux to Na^+^-influx at high extracellular Na^+^-concentrations and membrane potentials clamped close to the reversal potential. The same study described a different ChR variant from *Wobblia lunata*, the *Wobblia* inhibitory channelrhodopsin WiChR, with an increased selectivity of K^+^ over Na^+^ compared to both *Hc*KCRs (Vierock et al. 2022). A recent study by Ott *et al*. tested light-gated KCRs, including variants of *Hc*KCR1 and -2, in different animal models, among which was the nematode *Caenorhabditis elegans*. They expressed the channels pan-neuronally and observed a decrease in the crawling speed of the animals upon illumination (Ott et al. 2024). Yet, since both hyperpolarization and strong depolarization of cholinergic neurons can cause macroscopic locomotion slowing, it is unclear how slowing was affected.

We thus investigated animals expressing the KCRs *Hc*KCR1 and WiChR in cholinergic neurons and/or body wall muscle (BWM) cells of *C. elegans*, using a body length assay that reports on both hyper- and depolarization (Bergs et al. 2018). Similar to what has been described by Ott et al., 2024, we observed a speed reduction of the animals upon light exposure. However, our data suggest that, during continuous illumination, especially at high light intensities, such a behavioral effect might not be caused by hyperpolarization but instead by overexcitation of the muscles. This indicates that, while KCRs can be used to effectively hyperpolarize excitable cells in *C. elegans* under certain experimental conditions, they will be less effective, or potentially even have a depolarizing effect, using different conditions.

## Results

K^+^-channels in *C. elegans* are, among other functions, involved in the regulation of action potentials (APs) that drive muscle contraction. Specifically, in BWM cells, voltage-gated K^+^-channels induce outward-directed currents required for AP termination (Gao and Zhen 2011; Liu et al. 2011). Previously established hyperpolarizing optogenetic tools, such as light-activated ion pumps or ACRs expressed in BWM cells, upon illumination, can induce muscle relaxation resulting in an increase of the body length of the animals (Zhang et al. 2007; Bergs et al. 2018). Hence, to confirm the ability of the green light-reactive *Hc*KCR1 to hyperpolarize excitable cells in *C. elegans*, the channel was expressed in cholinergic neurons and the effect of its activation on the mean body length of the animals was investigated.

Initial experiments investigating the body length change of different transgenic strains expressing *Hc*KCR1 in cholinergic neurons (**Fig. 1A, B**) revealed unexpected results: Three of four different transgenic lines displayed a reduction in body length after stimulation of the channel at high light intensities (**Supplemental Fig. S1**; 535 nm, 4.5 mW/mm^2^). This would be expected for excitation and contraction of muscle cells, and indicates depolarizing effects, while hyperpolarization and thus inhibition would cause elongation of the animals. A fourth strain displayed a different phenotype, as the animals elongated briefly before returning to the initial length. After termination of the light pulse, contraction was observed, which may be attributed to post-inhibitory rebound, an effect also observed for other inhibitory tools in *C. elegans*, such as the chloride channel ACR2 in BWM cells (Bergs et al. 2018). Due to the brief apparent hyperpolarizing effect, this strain was selected for further experiments. Continuous light was applied for 10 s at 1 and at 0.1 mW/mm^2^, which is comparable to what Ott *et al*. used for *Hc*KCR1 activation in *C. elegans* neurons (Ott et al. 2024). For both conditions, the animals ceased moving during illumination (**Supplemental Video S1**), in agreement with the findings of the previous study. While high-intensity illumination largely induced only a brief hyperpolarizing phase despite ongoing illumination, low-intensity stimulation allowed achieving stable elongation throughout, suggesting that these conditions provide a more uniform hyperpolarization (**Fig. 1C, D**). No significant changes in body length were observed for control animals that were not supplemented with all-*trans* retinal (ATR), the ChR chromophore.

**Fig. 1:**
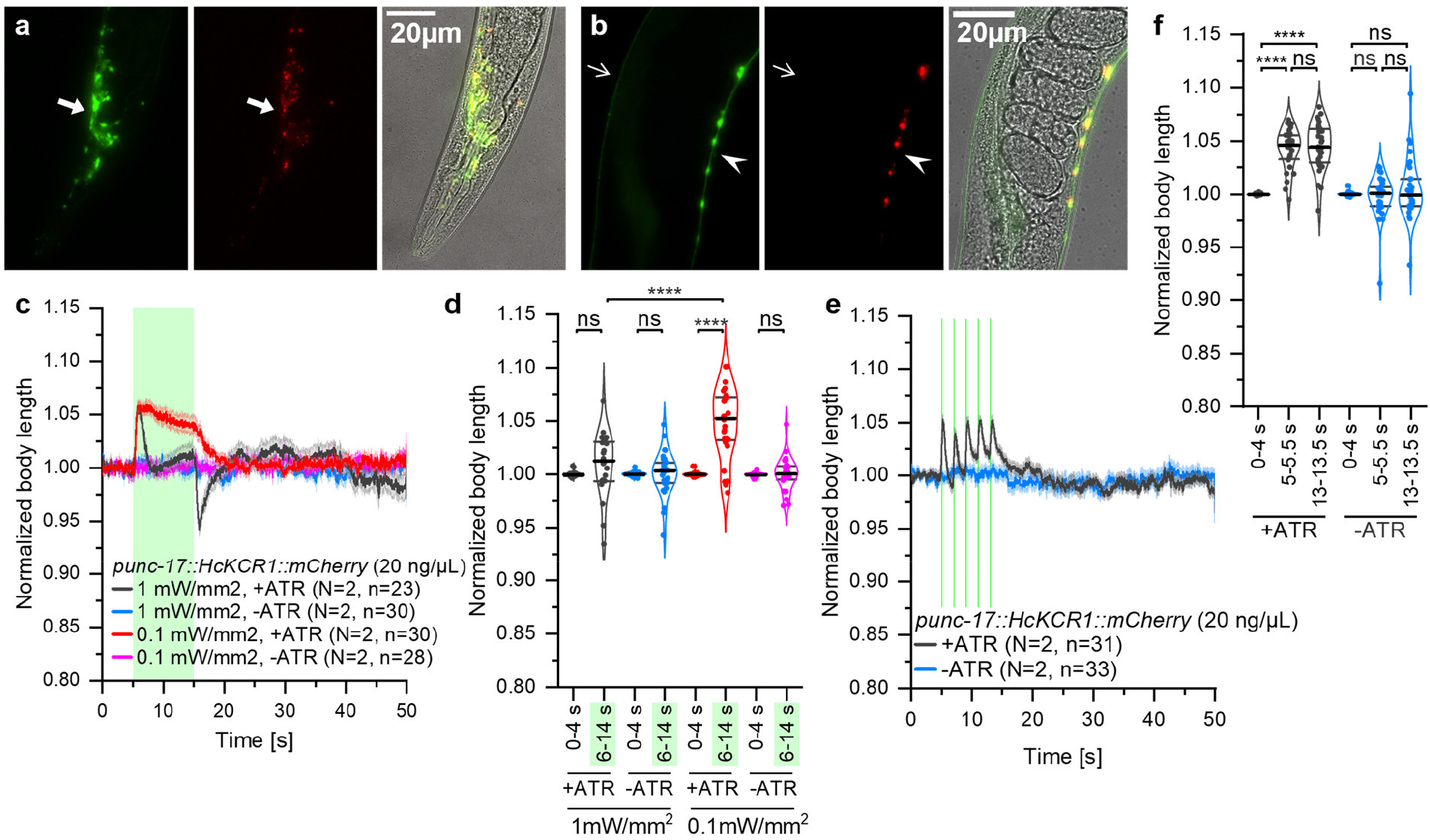
*Hc*KCR1 can be used to stably hyperpolarize cells at low light intensities. **(A, B)** Fluorescence micrographs of animals expressing p*unc-17::HcKCR1::mCherry::SL2::GFP* (20 ng/μL) in cholinergic neurons. Depicted are images of GFP fluorescence (left), mCherry fluorescence (middle) and overlays with transmission light images (right), showing the nerve ring (**A**, indicated by arrows) or the ventral and dorsal nerve cords (**B**, indicated by arrow heads and open arrows, respectively). **(C)** Body length of animals expressing p*unc-17::HcKCR1::mCherry::SL2::GFP* (20 ng/μL), grown on OP50 with or without ATR, normalized to the mean value of seconds 0 to 4. 10 s of continuous light stimulation with 535 nm light at an intensity of 0.1 or 1 mW/mm_2_ are indicated as a green shade. **(D)** Statistical analysis of the relative body lengths of single animals depicted in **C. (E)** As in **C**, but with pulsed light stimulation with a duration of 100 ms and a frequency of 0.5 Hz at an intensity of 1 mW/mm^2^. **(F)** Statistical analysis of the relative body lengths of single animals depicted in **E. (C, E)** Shown are mean ± SEM. N = number of individual experiments, n = total number of animals. **(D, F)** Median, 1^st^ and 3^rd^ quartile are indicated. Mixed-effect model analysis (REML) with Šidák’s multiple comparisons test was performed. ns, not significant (p>0.05), **** p<0.0001.

We next determined whether *Hc*KCR1 can be used as a reliable hyperpolarizing tool also while applying higher light intensities, by using pulsed stimulation (**Fig. 1E, F**). Applying five green light pulses (1 mW/mm^2^, 100 ms, 0.5 Hz) caused repeated elongation (i.e., hyperpolarization) and prevented the overshoot reaction observed after continuous illumination at the same intensity. The increased body length induced by the 5^th^ pulse was not significantly different from that of the 1^st^ one, suggesting that optimization of pulse length and frequency might also allow stable hyperpolarization under high intensity stimulation.

The return to initial body length despite ongoing illumination could indicate fast desensitization of *Hc*KCR1, however, this is not expected based on previous publications (Govorunova et al. 2022a; Ott et al. 2024). Given that most transgenic lines exhibited depolarizing effects, and since *Hc*KCR1 was described to have residual Na^+^-conductivity, we reasoned that there could be a change in the conductance during the prolonged activation, and resulting Na^+^-influx might counteract the hyperpolarizing effects of K^+^-efflux. Alternatively, conductance of the channel might be high enough to alter the K^+^ gradient sufficiently to prevent further K^+^-flux. WiChR was described to have superior selectivity for K^+^ over Na^+^ (Vierock et al. 2022), thus we next expressed WiChR in excitable cells, and performed similar experiments.

First, we tested animals expressing WiChR in cholinergic neurons (**Fig. 2A**). The expression pattern was similar to what was observed for *Hc*KCR1 (**Fig. 1A, B**). We then performed body length analyses before, during and after illumination. Upon continuous illumination with high- (1 mW/mm^2^) or low-intensity (0.1 mW/mm^2^) blue light, animals displayed only a very small, non-significant increase in body length, but instead contracted right after the beginning of the light pulse (**Fig. 2B, C**). This is in agreement with the contraction of muscle cells induced by the release of acetylcholine, indicating predominant activation (depolarization) of the cholinergic neurons. As described for worms expressing *Hc*KCR1 pan-neuronally (Ott et al. 2024), stimulation with lower intensities led to a drop in the crawling speed of the animals (measured with a multiworm tracker; Swierczek et al.. 2011; **Fig. 2D, E**). Surprisingly, the animal speed did not return to initial levels after the light pulse terminated; instead the animals remained immobile for ca. 1 minute, before slowly increasing their speed again over the next 3 minutes. We examined the behavior of individuals and found that the illumination induced no or only a brief body relaxation, but mainly contraction, which was followed by coiling behavior (**Supplemental Video S2**). This is similar to what we previously observed during prolonged depolarization of cholinergic neurons using the Na^+^-conducting ChR2 (Liewald et al. 2008), and is due to circuit effects in the motor neuron circuit, involving concomitant activation of GABAergic motor neurons. These findings, along with the reduced body length (**Fig. 2B, C**) show that WiChR activation induces predominantly depolarization rather than hyperpolarization, under the conditions tested, and that slowing of locomotion is due to overexcitation of the motor neurons. We also tested whether pulsed illumination of WiChR would allow more obvious hyperpolarization (**Fig. 2F, G**). While the 1^st^ pulse caused some, yet non-significant elongation, again depolarizing effects dominated, though they were slowed in their onset and the contraction less pronounced than for continuous illumination.

**Fig. 2:**
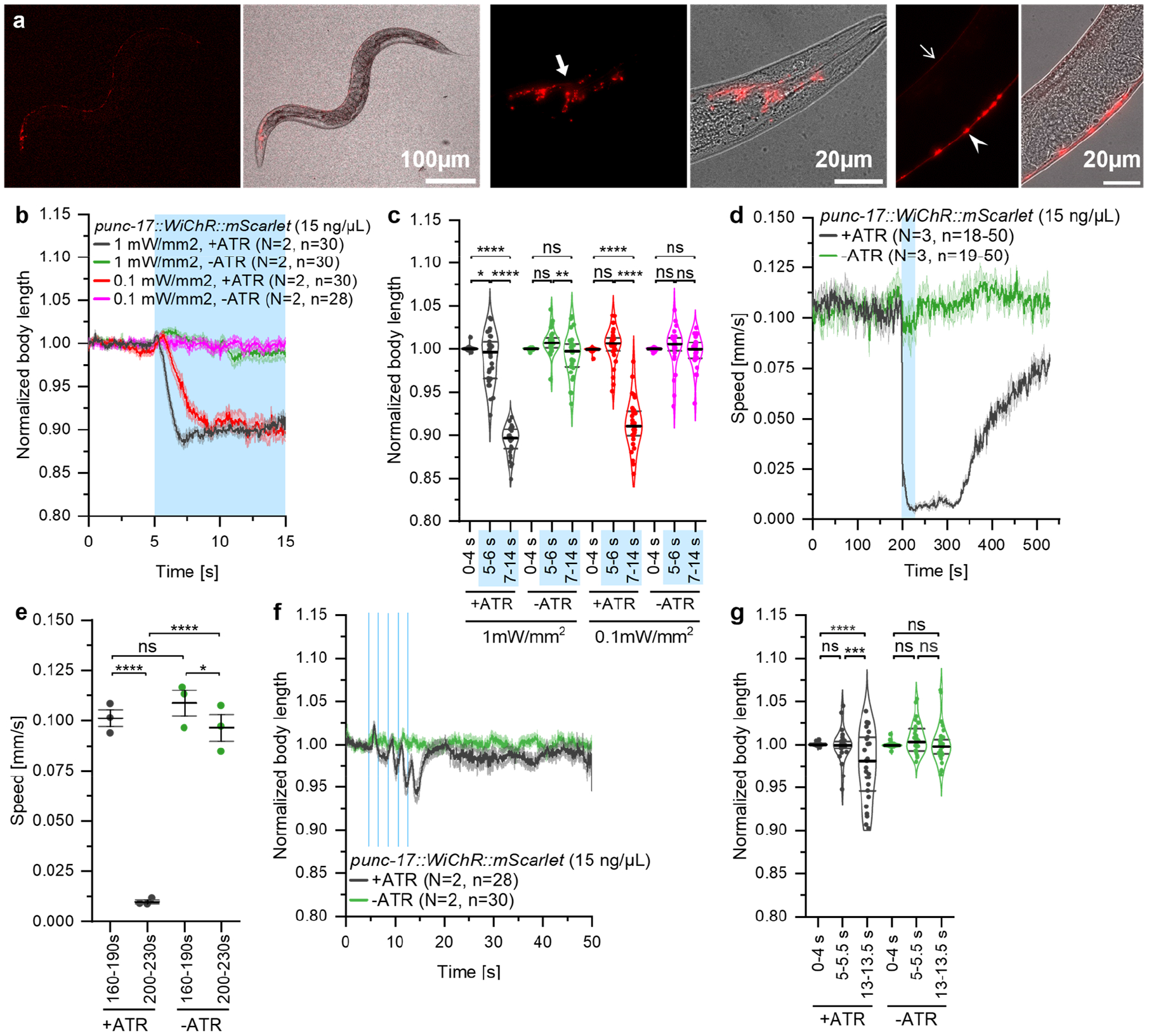
Speed decrease caused by WiChR activation in cholinergic neurons is induced by overexcitation rather than inhibition. **(A)** Fluorescent micrographs of animals expressing *punc-17::WiChR::mScarlet* (15 ng/μL) in cholinergic neurons. Depicted are fluorescence images (top) and overlays with transmission light images (bottom), showing an entire animal (left), the nerve ring (middle, indicated by an arrow) or the ventral and dorsal nerve cords (right, indicated by arrowhead and open arrow, respectively). **(B)** Body length of animals expressing *punc-17::WiChR::mScarlet* (15 ng/μL), grown on OP50 with or without ATR, normalized to the mean value of seconds 0 to 4. 10 s of continuous light stimulation with 470 nm light at an intensity of 0.1 or 1 mW/mm^2^ are indicated as a blue shade. **(C)** Statistical analysis of the relative body lengths of single animals depicted in **B. (D)** Off-food crawling speed of animals expressing *punc-17::WiChR::mScarlet* (15 ng/μL), grown on OP50 with or without ATR. 30 s of blue light stimulation with 470 nm light at an intensity of 0.1 mW/mm^2^ are indicated as a blue background. **(E)** Statistical analysis of the crawling speeds depicted in **D**. Mean values ± SEM are indicated. **(F)** As in **B**, but with pulsed light stimulation (100 ms pulses, 0.5 Hz, 1 mW/mm^2)^. **(G)** Statistical analysis of the relative body lengths of single animals depicted in **F. (B, D, F)** Shown are mean ± SEM. **(B, F)** N = number of individual experiments, n = total number of animals. **(D)** N = number of individual experiments, n = number of animals per experiment. **(C, G)** Median, 1^st^ and 3^rd^ quartile are indicated. **(C, E, G)** RM two-way ANOVA (**C, E**) or mixed-effect model analysis (REML) (**G**) with Šidák’s multiple comparisons test was performed. ns, not significant (p>0.05), * p<0.05, ** p<0.01, *** p<0.001, **** p<0.0001.

Thus far, we tested evoked effects of KCRs in cholinergic neurons, using indirect read-outs (behavior, muscle contraction). However, when expressed in BWM cells, ChRs can affect muscle activity directly, providing a more immediate assay for the nature of the evoked photocurrents. We thus expressed WiChR in BWM cells. Fluorescence microscopy confirmed the presence of mScarlet-labelled WiChR protein at the plasma membrane of BWM cells (**Fig. 3A**). Next, WiChR activation was induced in individual animals through 10 s of continuous 1 or 0.1 mW/mm^2^ intensity 470 nm light, which caused paralysis in worms supplemented with ATR (**Supplemental Video S3**). Analysis of the body length revealed only an insignificant brief elongation of the ATR-supplemented worms for 1 mW/mm^2^, followed by a strong contraction that only slowly subsided after the end of the light pulse (**Fig. 3B, C**). This confirmed the suspected reversal of currents upon continuing illumination and resembles what we found for most of the transgenic strains expressing *Hc*KCR1 (**Supplemental Fig. S1**). The reduction of the light intensity caused slightly different effects in that a first, brief, but significant hyperpolarization could be achieved (**Fig. 3B, C**). Yet, also the 0.1 mW/mm^2^ stimulation did not prevent the subsequent depolarization. Since, compared to HcKCR1, WiChR is more light-sensitive (Vierock et al. 2022), it would potentially require even lower light intensities for the same effect. We also tested application of pulsed light (1 mW/mm^2^, 100 ms, 0.5 Hz; **Fig. 3D, E**). The 1^st^ pulse induced a significant hyperpolarization in ATR-supplemented animals, as their body lengths significantly increased compared to prior to stimulation. However, the mean amplitude of the effect appeared to decrease with each pulse, and the length after the 5^th^ pulse was decreased compared to the initial length. Nevertheless, the duration of the elongation was increased compared to the effects of continuous stimulation, and the amplitude of the contraction was decreased. Thus, by optimizing the frequency and duration of light pulses, one may extend the duration of the hyperpolarization.

**Fig. 3:**
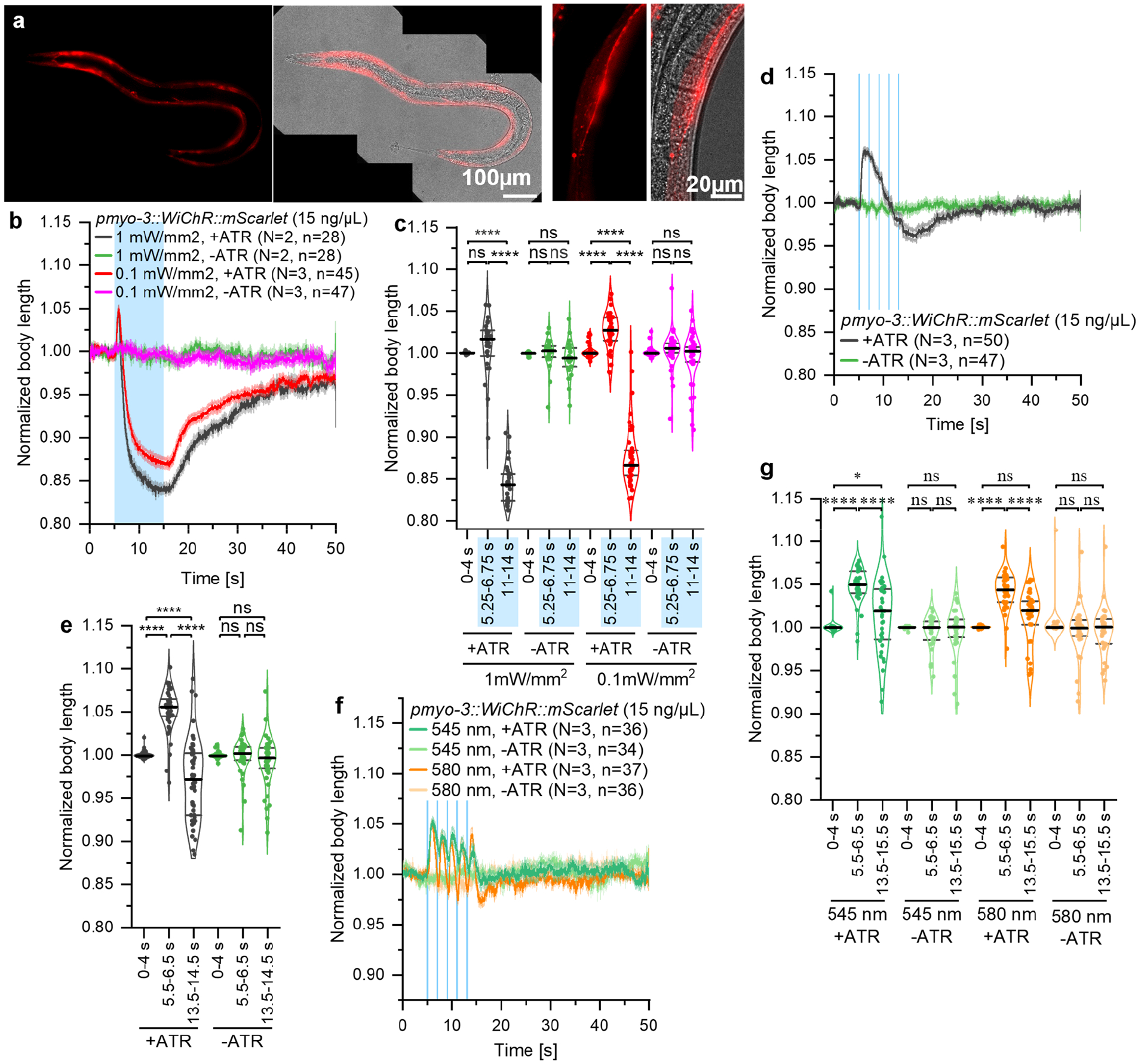
WiChR activation in BWM cells leads to a brief hyperpolarization followed by a continuing depolarization. **(A)** Fluorescent micrographs of BWM cells of animals expressing *pymo-3::WiChR::mScarlet* (15 ng/μL). Depicted are fluorescence images (left) and overlays with transmission light images (right), showing an entire animal (left images) or a single muscle cell (right images). **(B)** Body length of animals expressing p*ymo-3::WiChR::mScarlet* (15 ng/μL), grown on OP50 with or without ATR, normalized to the mean value of seconds 0 to 4. 10 s of continuous light stimulation with 470 nm light at an intensity of 0.1 or 1 mW/mm^2^ are indicated as a blue shade. **(C)** Statistical analysis of the relative body lengths of single animals depicted in **B. (D)** As in **B**, but with pulsed light stimulation (100 ms, 0.5 Hz, 470 nm, 1 mW/mm^2^). **(E)** Statistical analysis of the relative body lengths of single animals depicted in **D. (F)** As in **D**, but using light of 545 or 580 nm. **(G)** as in **E**, for animals depicted in **D. (B, D, F)** Shown are mean ± SEM. N = number of individual experiments, n = total number of animals. **(C, E, G)** Median, 1^st^ and 3^rd^ quartile. Mixed-effect model analysis with Šidák’s multiple comparisons test was performed. ns, not significant (p>0.05), **** p<0.0001.

Last, we tested whether illumination at different wavelengths, i.e. avoiding the peak of the WiChR action spectrum, may further reduce channel activation and allow shifting the effects to the hyperpolarizing regime. Accordingly, when we expressed WiChR in ND7/23 cells and measured photoevoked currents, we observed, close to the reversal potential, a late photocurrent reversion during long illumination with 480 nm light, that could be avoided with violet or orange illumination (**Supplemental Fig. S3**). We thus compared effects evoked in animals expressing WiChR in BWM cells using continuous or pulsed illumination with 470, 545 and 580 nm light, at different intensities (**Fig. 3F, G; Supplemental Fig. S2**). These experiments showed that low intensity, continuous illumination with 545 nm light had a more pronounced hyperpolarizing effect than high-intensity illumination, but still the net effect of the WiChR mediated currents inversed to depolarization (**Supplemental Fig. S2**). When using pulsed stimulation, both at 545 and 580 nm, we could effectively prevent the depolarizing effects, with the body length oscillating between initial, and elongated amplitudes (**Fig. 3F, G**).

## Discussion

Here, we compared and benchmarked the potential of hyperpolarizing KCRs, *Hc*KCR1 and WiChR, for use in inhibition of excitable cells in *C. elegans*. The presented data suggests that, depending on the conditions of the illumination, and likely, expression levels, KCRs are able to hyperpolarize *C. elegans* excitable cells. However, they also highlight that these proteins can induce depolarization, with WiChR potentially having a higher tendency to do so than *Hc*KCR1. Yet, also for *Hc*KCR1, we have observed transgenic strains that induced a similar, depolarizing effect in cholinergic neurons. These findings suggest that the observed differences might not only be due to intrinsic differences in the characteristics of the channels but may also be influenced by likely different expression levels, or other factors that are difficult to control during the generation of transgenic strains, emphasizing the need to test different lines. By alteration of light intensity, color and duration of illumination, the net outcome of hyperpolarization *vs*. depolarization can be shifted to predominant hyperpolarization.

The reason for the switch from hyper- to depolarization is currently unclear. Although, this effect resembles the reversal of currents that has been described for *Hc*KCR1 at high extracellular Na^+^-concentrations and membrane potentials close to the reversal potential, the same effect was even more pronounced for WiChR in *C. elegans* muscles or neurons, that – in patch clamp recording on mammalian cells - showed less inactivation and more negatively shifted reversal potentials (Vierock et al. 2022). Subsequent changes in K^+^-selectivity had also been reported for the slow *Hc*KCR1_C110A mutant and correlated with three distinct blue shifted photocycle intermediates (Sineshchekov et al. 2024). These were associated with alternative photocycles and might represent different conductive states of the channel that are populated dependent on illumination time and intensity similar as observed in other ChRs before (Kuhne et al. 2019). And although a comparable analysis for alternative photocycle reactions for *Hc*KCR1 and WiChR is still missing, different conductive states with partly reduced K^+^ selectivity could partly explain the observed phenotypes.

Noteworthy however, recovery of depolarization in *C. elegans* was slow and surpassed the expected channel closure by far. This indicates that the biophysical properties of both channels alone might not be sufficient to explain the observed change of membrane voltage manipulation. Thus, potentially, some unknown factors in the cellular environment of *C. elegans* may additionally promote this effect and enable such a reversal even at physiologic membrane potentials. One possible explanation for the observed effects might be found in the large conductance mediated by the KCRs, estimated as 700 fS for *Hc*KCR1 by noise analysis (Govorunova et al. 2022a), and potentially even larger for WiChR, because of reduced inactivation during continuous light. Other ChRs exhibit a unitary conductance that is orders of magnitude smaller (Feldbauer et al. 2009; Zerche et al. 2023). Should the K^+^-current be large enough, a macroscopic change in the K^+^-gradient might occur that would lead to seizing of K^+^-currents, already during the illumination. Once no K^+^-efflux occurs, the residual Na^+^-conductivity could enable an influx of Na^+^, thus leading to depolarization. The long-lasting effects could be explained if one considers that the reinstatement of the native, physiological Na^+^/K^+^-gradient, which needs to be achieved by the Na^+^/K^+^-ATPase, and which may require a considerable time, as indicated by the long-lasting depolarizing and locomotion slowing effects following KCR activation in cholinergic neurons. It is currently unknown whether this phenomenon is specific for the nematode, possibly induced by interactions with native proteins, or by the physiological properties of the Na^+^/K^+^-ATPase, or if it is also observed in other organisms. The phenotypes are in some aspects contradictory to the characteristics of *Hc*KCRs in *Drosophila*, as it has been found by Ott *et al*. that higher light intensities increase the inhibitory potential of the channels (Ott et al. 2024). The intensities used in these experiments, however, were 2- to 4-fold lower than those for the low-intensity measurements we described here.

In sum, our results stress the requirement to explore the mechanisms underlying behavioral responses, such as a decrease in crawling speed. Further, they emphasize the need for testing different transgeninc lines and experimental conditions when *Hc*KCR1 or WiChR are to be used for their hyperpolarizing properties. For *C. elegans*, if the experimental design and setup allow using a green light-activated inhibitory tool, HcKCR1 might be preferable as its application was more straightforward and it allowed more stable hyperpolarization for a longer duration. Here, future applications might also consider the *Hc*KCR1-H225F (Kali-1) or *Hc*KCR1-C29D mutants that were also reported to improve K^+^-selectivity of the channel (Tajima et al. 2023, Vierock et al. 2022), and for *Hc*KCR1-C29D also reduced changes in K^+^-selective conductance during continuous illumination (Morizumi et al. 2023). Finally, however, irrespective of the exact KCR used, both intensity and illumination pattern must be optimized, and the behavior of the channels under these conditions confirmed, to ensure that the desired effect, i.e. inhibition, can be achieved.

## Acknowledgements

We thank Katharina Kuhlmeier, Franziska Baumbach and Olivia Herczynski for expert technical assistance. We are indebted to Elena Govorunova and John Spudich for sharing of information prior to publication of their initial KCR paper, and to members of the Gottschalk lab for fruitful discussions.

## Funding

This work was funded by the Deutsche Forschungsgemeinschaft (DFG), grants SFB1080-B2 to AG, and EXC-2049–390688087 and SFB1315 to JV, as well as by resources from Goethe University to AG.

## Conflict of Interest

The authors declare no competing financial interest

## Materials & Methods

### Molecular biology, *C. elegans* maintenance & transgenic animals

The DNA construct for expression of HcKCR1 in cholinergic neurons was generated by three-fragment Gibson assembly of the HcKCR1 coding sequence, a fragment encoding for mCherry::SL2::GFP, amplified from pXY12 (*pdat-1::ChR2::mCherry::SL2::GFP*) and the vector backbone of pMSE01 (*punc-17::Chrimson (S169A)*).

The WiChR-mScarlet coding sequence was cloned for expression in BWMs through digest of pJV888 (*pCMV::WiChR::mScarlet*) with EcoRI and NheI and ligation into the vector backbone of pFB011 (*pmyo-3::ChR-XXM-2*.*0::mVenus*). For expression in cholinergic neurons, pJV855 was digested with EcoRV and NheI and the coding sequence ligated into RM#348p (*punc-17::unc-54-3’UTR*).

Animals were grown at room temperature on nematode growth medium (NGM) plates seeded with *E. coli* OP50-1. Transgenic strains were generated by microinjection of 15 or 20 ng/µL plasmid DNA. For the strains expressing WiChR, 1.5 ng/µL pmyo-2:CFP was co-injected as a marker for transgene expression.

The following strains were generated and used: **ZX3283**: wild type; *zxEx1394[punc-17::HcKCR1::mCherry::SL2::GFP]*, **ZX3706**: N2*;zxEx1472[pmyo-3::WiChR::mScarlet; pmyo-2::CFP]*, **ZX3888**: *lite-1(ce314) X*.*;zxEx1519[punc-17::WiChR::mScarlet; pmyo-2::CFP]*.

### Fluorescence imaging

Fluorescence imaging was performed on a Zeiss Axio Observer Z1 using 10x or 40x objectives. Young adult animals were placed on imaging pads made of 7% agarose (Carl Roth) in M9 medium in approx. 5 μL of 20 mM tetramisole hydrochloride (Sigma-Aldrich) in M9. Images were processed using FIJI (ImageJ), and partially processed using the “Image Stitching” plugin (Preibisch et al. 2009).

### Body length measurements

Transgenic L4 larvae were placed on NGM plates seeded with OP50 bacteria with or without 200 μM ATR (Sigma Aldrich). The plates were wrapped in aluminum foil and the worms grown at room temperature overnight.

On the measurement days, single animals were moved to fresh unseeded plates which were mounted on a Zeiss Axioscope A1 microscope, equipped with a Canon Powershot G9 camera. During the assays, animals were illuminated with light from a 50 W HBO lamp using a 690/50 emission filter and 470/40 or 535/30 bandpass filters for excitation of WiChR and HcKCR1, respectively. The light intensity was adjusted to approx. 0.1 mW/mm^2^ or 1 mW/mm^2^, as indicated. For measurements with continuous illumination, animals were observed for 5 s, then continuously illuminated for 10 s. For pulsed illumination, five pulses were applied with a duration of 100 ms and a frequency of 0.5 Hz after 5 s. Following the light pulses, the recordings were continued for 35 s. Simultaneous start of the video acquisition and illumination pattern was controlled by an Arduino. All videos were analyzed using the WormRuler software (version 1.3) (Seidenthal et al. 2022). The mean body lengths over the first 4 s of the measurements were used for normalization. Due to the inability of the program to measure the body lengths of coiled animals, for videos of animals expressing WiChR in cholinergic neurons, only the first 15 s were used for further analyses.

### Crawling speed measurements

Transgenic L4 animals were prepared as described for the body length measurements. The following day, the worms were flushed off the plates and washed three times using M9 before removing the majority of the liquid and pipetting them onto unseeded NGM plates. The remaining M9 buffer was allowed to dry and subsequently, to avoid any pre-stimulation, the animals were kept in the dark for 15 min before the measurement was started.

Crawling assays were performed using a multiworm tracker setup (MWT) (Swierczek et al. 2011) composed of a camera (Falcon 4M30, DALSA), an infrared transmission light source (6 WEPIR3-S1, WINGER, 850 nm 3 W) and an LED ring (Alustar 3 W 30°, ledxon, 470 nm) allowing stimulation of NGM plates placed in the setup with blue light. The worms were observed for 5 min without stimulation, followed by illumination at a wavelength of 470 nm and an intensity of approx. 0.1 mW/mm^2^ for 30 s and 5 min observation after the light stimulus. The crawling speed was recorded using the “Multi-Worm Tracker” software (version 1.3.0) and the tracks were extracted using “Choreography”. We have observed previously that the software does not properly track the animals immediately after starting the assay, so the first 100 s of the recordings were discarded.

### Molecular Biology, Cell Culture and Whole-Cell patch clamp with ND7/23 cells

Mammalian-codon optimized HcKCR1 (GenBank MZ826861) and WiChR (Addgene #195190/GenBank OP710241) were expressed in rodent neuroblastoma cells similar as previous described (Vierock et al. 2022). Accordingly, ND7/23 cells (ECACC 92090903 from Sigma Aldrich) were seeded on polylysin coated coverslips at a concentration of 0.8 × 105 cells/ml and cultured at 5% CO2 and 37°C in Dulbecco’s minimal essential medium supplemented with 5% fetal bovine serum, 100 μg/ml penicillin/streptomycin and 1 μM all-*trans*-retinal (all from Merck KGaA, Darmstadt, DE) and subsequently transfected using the FuGENE HD Transfection Reagent (Promega, Madison, US) 28–48 h before measurement.

Whole-cell voltage clamp recordings were performed using an Axopatch 200B amplifier, filtered at 2 kHz, digitized at 10 kHz (DigiData 1550B) and acquired using Clampex 10.7 Software (all from Molecular Devices, San Jose, US). Patch pipettes were prepared using a P-1000 micropipette puller (Sutter Instruments, Novato, US) from borosilicate glass capillaries (GB150F-8P; Science Products GmbH, Hofheim, DE) with a resistance of 1.5 to 2.5 MΩ. A reference AgCl-electrode was connected to the bath solution with a 140 mM NaCl 1.5% agar bridge. The pipette solution contained 110 mM potassium-D-gluconate, 1 mM NaCl, 2 mM CaCl2, 2 mM MgCl2, 10 mM EGTA and 10 mM HEPES at pHi 7.2 (glucose was added up to 290 mOsm) and the bath solution contained 110 mM NaCl, 1 mM KCl, 2 mM CaCl_2_, 2 mM MgCl_2_, and 10 mM HEPES at pHe 7.2 (with glucose added up to 310 mOsm). Light used for excitation was provided by an OptoScan Monochromator (Cairn Research, Kent, UK) with an emission bandwidth of 10 nm and intensities adjusted to gain equal photon densities for all wavelengths with 480 nm having an intensity of 0.81 mW/mm^2^ and 520 nm of 0.75 mW/mm^2^. The light source was coupled into an Olympus IX70 inverted microscope (Olympus, Tokyo, JP) with a LUMPlanFLN 60x water objective (Olympus, Tokyo, JP) and a 90/10 beamsplitter (BSX10R; ThorLabs, Newton, US) and controlled using a mechanical shutter (UniBlitz VS25; Vincent Associates, Rochester, US). Action spectra were recorded with 15 s illumination at -80 mV for WiChR or -75 mV HcKCR1. Measurements were analyzed using Clampfit 10.7 software (Molecular Devices, San Jose, US), Microsoft Excel and GraphPad Prism 9.5.1 (GraphPad Software, Boston, US). Photocurrent traces were baseline corrected, filtered and reduced in size for displaying purposes.

## Supplemental Figures

**Supplemental Figure S1:**
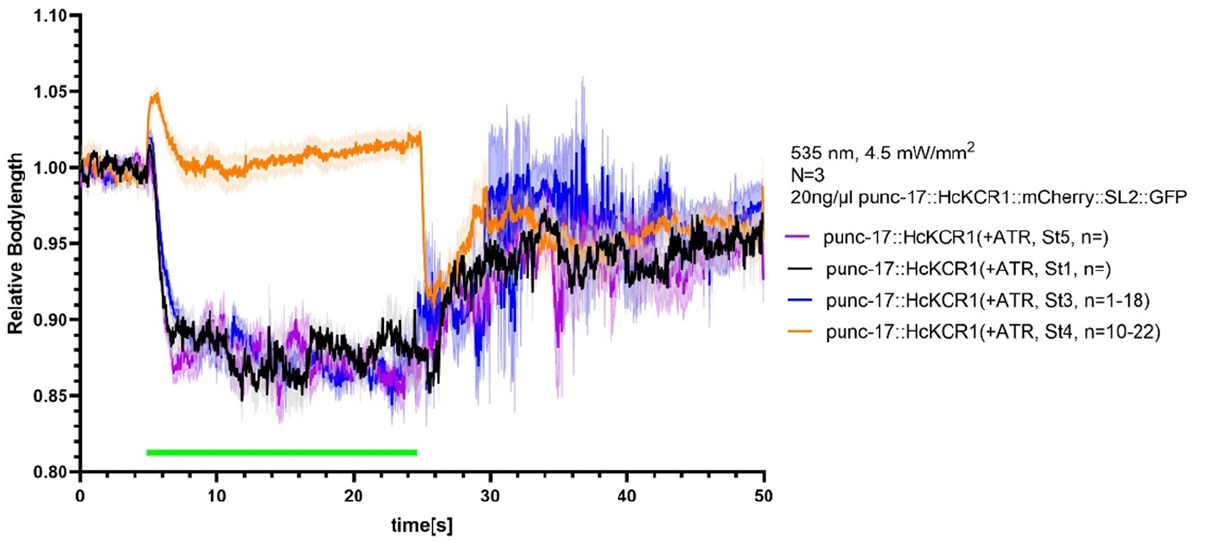
HcKCR1 can induce a brief hyperpolarization or an intense, continuous depolarization. Body length of animals expressing *punc-17::HcKCR1::mCherry::SL2::GFP* (20 ng/μL), grown on plates with ATR, normalized to the mean value of seconds 0 to 5. 20 s of continuous light stimulation with 535 nm light at an intensity of 4.5 mW/mm^2^ are indicated by a green line.

**Supplemental Figure S2:**
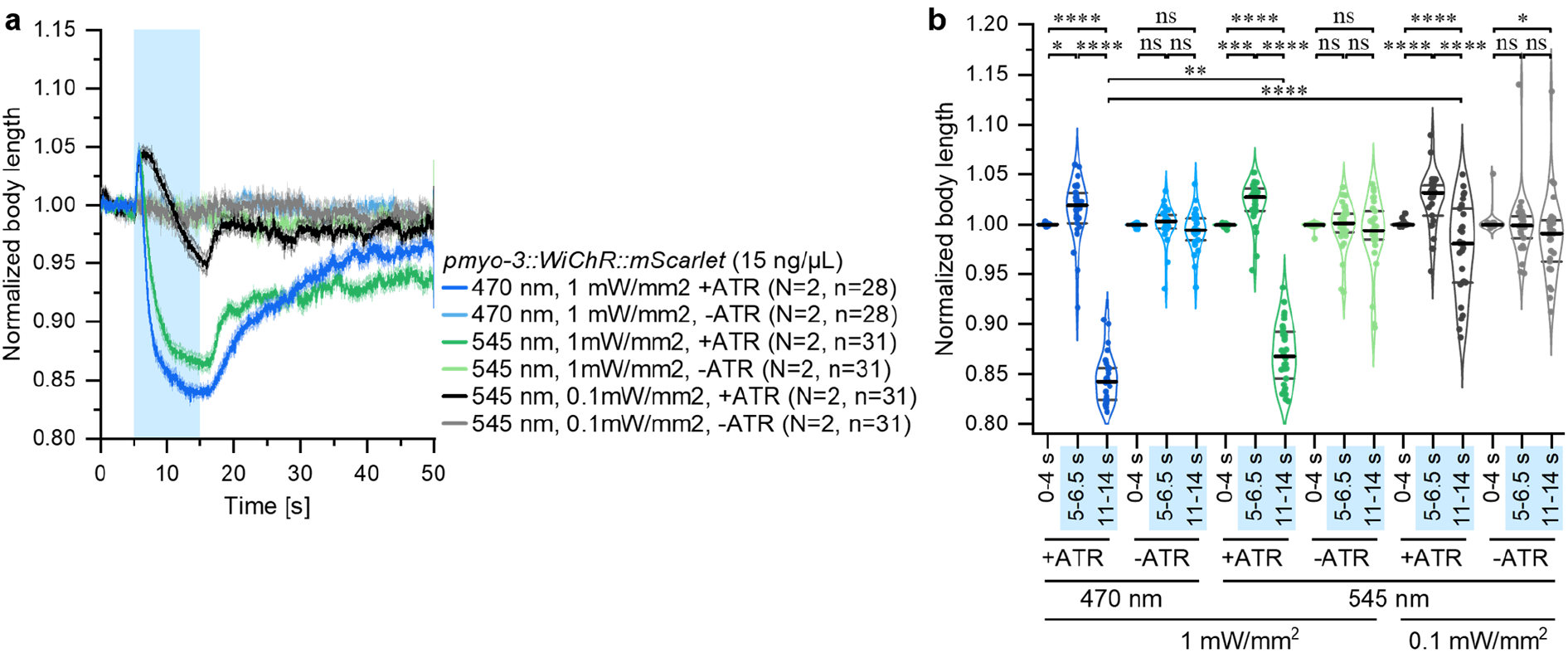
WiChR-induced depolarization can be reduced by optimizing wavelength and intensity. **(A)** Body length of animals expressing *pymo-3::WiChR::mScarlet* (15 ng/μL), grown on plates with or without ATR, normalized to the mean value of seconds 0 to 4. 10 s of continuous light stimulation with 470 nm or 545 nm light at an intensity of 0.1 or 1 mW/mm^2^ are indicated as a blue shade. The data shown for illumination with 470 nm light at an intensity of 1 mW/mm^2^ (+ and -ATR) is the same that is depicted in **Fig. 2 B**. Shown is mean ± SEM. N = number of individual experiments, n = total number of animals. **(B)** Statistical analysis of the relative body lengths of single animals depicted in **A**. Median, 1^st^ and 3^rd^ quartile are indicated. Mixed-effect model analysis (REML) with Šidák’s multiple comparisons test was performed. ns, not significant (p>0.05), * p<0.05, ** p<0.01, *** p>0.001, **** p<0.0001.

**Supplemental Figure S3:**
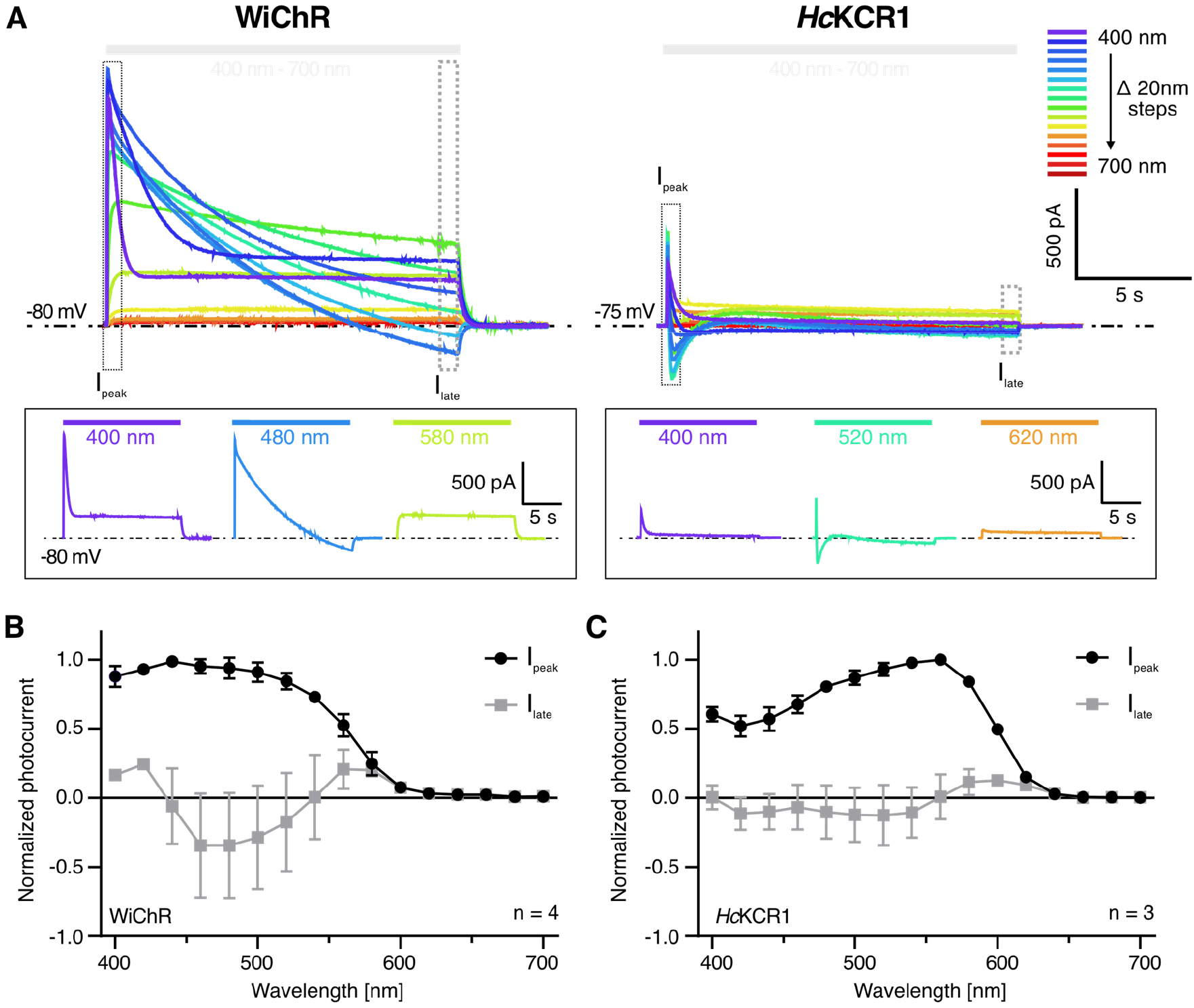
Wavelength-dependent photocurrent reversion of KCRs with prolonged illumination in ND7/23 cells. (**A**) Representative photocurrent traces of WiChR at -80 mV and of *Hc*KCR1 at -75 mV with 15 s illumination at wavelengths from 400 nm to 700 nm and equal photon count. Measuring solutions contained 110 mM Na_e_^+^ Cl / 1 mM K_e_^+^Cl in the extracellular buffer and 110 mM K_i_^+^ gluconate / 1 mM Na_i_^+^ Cl in the patch pipette - both at pH_e/i_ 7.2. Insets at the bottom highlight specific illumination conditions for both channels. Although initially exclusively outward directed photocurrents of both channels were observed, a change in direction during continuous illumination occurred at their absorption maximum with kinetics that depended on the channel and that – especially for WiChR - varied importantly for different cells. Surprisingly however, no photocurrent inversion was observed for violet- and yellow or orange light. The resulting spectra of the peak and the late photocurrent of (**B**) WiChR (n=4) and (**C**) *Hc*KCR1 (n=3) are normalized to peak photocurrents at λ_max_. Plots show mean ± SD.

